# Pallidal deep brain stimulation alters cortico-striatal synaptic communication in dystonic hamsters

**DOI:** 10.1101/2020.10.25.353847

**Authors:** Marco Heerdegen, Monique Zwar, Denise Franz, Valentin Neubert, Franz Plocksties, Christoph Niemann, Dirk Timmermann, Christian Bahls, Ursula van Rienen, Maria Paap, Stefanie Perl, Annika Lüttig, Angelika Richter, Rüdiger Köhling

**Affiliations:** Oscar Langendorff Institute of Physiology, Rostock University Medical Center; Institute of Applied Microelectronics and Computer Engineering, Faculty of Computer Science and Electrical Engineering, University of Rostock; Institute of General Electrical Engineering, Faculty of Computer Science and Electrical Engineering, University of Rostock; Institute of Pharmacology, Pharmacy und Toxicology, Faculty of Veterinary Medicine, University of Leipzig; Department Life, Light & Matter, University of Rostock; Department of Ageing of Individuals and Society, University of Rostock

**Keywords:** deep brain stimulation, globus pallidus, dystonia, *dt^sz^* hamster, inhibition

## Abstract

**Background:** Deep brain stimulation (DBS) of the globus pallidus internus (GPi) is considered to be the most relevant therapeutic option for patients with severe dystonias, which are thought to arise from a disturbance in striatal control of the GPi, possibly resulting in thalamic disinhibition. The mechanisms of GPi-DBS are far from understood. Hypotheses range from an overall silencing of target nuclei (due to e.g. depolarisation block), via differential alterations in thalamic firing, to disruption of oscillatory activity in the β-range. Although a disturbance of striatal function is thought to play a key role in dystonia, the effects of DBS on cortico-striatal function are unknown.

**Objective:** We hypothesised that DBS, via axonal backfiring, or indirectly via thalamic and cortical coupling, alters striatal network function. We aimed to test this hypothesis in the *dt^sz^*-hamster, an animal model of inherited generalised, paroxysmal dystonia.

**Methods:** Hamsters (*dt^sz^*-dystonic and non-dystonic controls) were bilaterally implanted with stimulation electrodes targeting the entopeduncular nucleus (EPN, equivalent of human GPi). DBS (130 Hz), and sham DBS, were performed in unanaesthetised animals for 3 hours. Synaptic cortico-striatal field potential responses, as well as miniature excitatory postsynaptic currents (mEPSC) and firing properties of medium spiny striatal neurons were subsequently recorded in brain slice preparations obtained from these animals immediately after EPN-DBS, to gauge synaptic responsiveness of cortico-striatal projections, their inhibitory control, and striatal neuronal excitability.

**Results:** DBS increased cortico-striatal responses in slices from control, but not dystonic animals. Inhibitory control of these responses, in turn, was differentially affected: DBS increased inhibitory control in dystonic, and decreased it in healthy tissue. A modulation of presynaptic mechanisms is likely involved, as mEPSC frequency was reduced strongly in dystonic, and less prominently in healthy tissues, while cellular properties of medium-spiny neurons remained unchanged.

**Conclusion:** DBS leads to dampening of cortico-striatal communication with restored inhibitory tone.

## Introduction

Primary dystonias were first recognised to be linked to brain, particularly to basal ganglia dysfunction by Charles Marsden and his group [1,2] only as late as in the 1970s. By now, it is being recognised that in dystonic patients, a pathological cortico-striatal function, and subsequent disturbance of striatal control of GPi, are likely to be important factors of dystonic dysfunction[1]. This disturbance is characterised by increased synaptic plasticity within the cortico-basal ganglia network [3–5], and speculated to result in a shift of the balance toward the direct pathway [6] (Fig. 5A). Beyond the basal ganglia, such network disturbances are reflected in loss of cortical inhibition [7–9], and a relative persistence of β-band synchronisation during movement initiation [10], as well as dominant low-frequency pallidal activity in the α-band at rest [11,12]. Indeed, a loss of inhibitory tone within the extended network is being discussed [13], although it is still unresolved whether the entire network or parts of it would be affected [14]. Although a contribution of cerebellar dysfunction is being assumed [5,15], an altered striatal function is likely a major causal factor in primary dystonias.

Deep brain stimulation (DBS) is clearly the most important innovation for the treatment of dystonias, and often the “only option for symptom reduction” [16]. As a consequence, clinical trials implementing DBS in dystonia [17,18] show that it is largely successful in patient groups particularly with idiopathic or genetic isolated dystonias. However, as much as the pathomechanisms of dystonia are still not fully understood, this knowledge gap extends even more so to the mechanisms underlying the effects of DBS in dystonia for several reasons: One is that DBS has most widely been investigated in patients with and animal models of Parkinson’s disease (PD) (see reviews [19,20]), which allows inference on DBS mechanisms in dystonia only in a limited way, also since the target nuclei (GPi vs. nucleus subthalamicus) are usually not the same. A second is that most of the hypotheses on this question are derived either from DBS in normal primates, or from DBS-like stimulation in vitro in normal rodent tissue, or from cortical or basal ganglia recordings of e.g. local fields in patients which obviously limit the extent to which the entire network can be assessed. Importantly, the effect of DBS in dystonias, in contrast to most motor symptoms of PD, require at least hours of stimulation, indicating that functional network changes likely occur [20]. To summarise the findings so far, the data from PD patients suggest that pathological oscillatory activity prominent in the β-band can be reduced by subthalamic DBS [11,21,22] – it is unknown whether this is the case for prominent resting α-band or transient β-band desynchronization during movement activity in dystonia. What is known is that pallidal DBS in dystonic patients does have network effects interpreted by the authors as inhibitory – increased cortical excitability and synaptic plasticity tested by e.g. paired associative stimulation or so-called cortical silent period using motor evoked potentials seem to be normalised [13,23–25] and firing of thalamic neurons is altered, albeit differentially (reduced in the majority of neurons, increased in a minority) [26]. It is thus safe to conclude that cortical excitability is somehow reduced by pallidal DBS, but nothing is known on alterations in the extended network, in particular regarding cortico-striatal functional connectivity. Looking at animal studies, in one investigation in normal primates, pallidal DBS completely silenced neuronal firing in this nucleus – presumably via activation of GABAergic afferents to the nucleus [27]. In contrast to this, a study on DBS-like stimulation in normal rat brain *in vitro* led to practically opposite effects, with high-frequency stimulation leading to prolonged afterdepolarisations mediated by cholinergic inputs, and no silencing of neurons [28] – thus the issue remains undecided. More importantly, animal studies so far were mainly conducted on healthy controls. In studies using animal models of dystonia, in turn, DBS-stimulation was delivered only under deep anaesthesia (urethane [29,30], or pentobarbital [31]). Even though anaesthesia (particularly urethane) is known to distort cortico-striatal connectivity [32], one of these studies does indicate that even DBS under pentobarbital anaesthesia does have a reducing effect on dystonia. Data on DBS effects in dystonia models in awake and behaving animals are completely lacking.

In view of the scant knowledge on the mechanisms of DBS, and in particular on excitability changes in the nuclei presumably being strongly involved in dystonic pathophysiology, i.e. the corpus striatum, we set out to test the lasting effect of prolonged (3 hours) pallidal DBS, presenting the first study so far conducting DBS in freely moving dystonic animals. For this, we chose an animal model which at least in many ways resembles the human situation of generalised paroxysmal dystonia, the *dt^sz^* hamster, which we have extensively characterised in the past [33–38]. Although the generalisability to human primary dystonia is unclear, there are important similarities: This strain shows spontaneous paroxysmal dystonic attacks which can also be provoked by handling and stress. As speculated for at least some human dystonias [3–5], this animal model is also associated with increased cortico-striatal excitability [39], on the basis of reduced intra-striatal GABAergic signalling, resulting in overall increased EPN/GPi inhibition [35,40]. Interestingly, this is in line with disturbed cortico-striatal communication [41] and increased pallidal inhibition also in DYT1 mouse [42]. As we could show in a recent study [43], short term DBS of the entopeducular nucleus with 130 Hz effectively reduces dystonic attacks. Importantly, in the present study we used the same DBS protocol in freely moving dt^sz^ hamsters to elucidate underlying mechanisms in electrophysiological studies on the cortico-striatal network. In this paper, we propose as a possible mechanism of DBS a dampening of cortico-striatal synaptic communication possibly due to presynaptic changes mediated anterogradely via thalamo-striatal or thalamo-cortical projections.

## Material and methods

### Animals

The experiments were carried out using two groups of age-matched dystonic *dt^sz^* mutant hamsters (inbred; total n=84), obtained by selective breeding (Institute of Pharmacology, University of Leipzig) as described previously [37], and two groups of age-matched non-dystonic control hamsters (Mesocricetus auratus, outbred, total of n=43) provided by a commercial breeder (JANVIER LABS; origin: Central Institute for Laboratory Animal Breeding, Hannover, Germany,). Dystonic *dt^sz^* hamsters display spontaneous dystonic attacks particularly after stress, as described below [37,38]. The animals were kept under controlled environmental conditions with a 14 h/10 h light/dark cycle and an ambient temperature of 23°C. Standard diet and water were supplied ad libitum.

After weaning at the age of 21 days, all groups of hamsters were screened for dystonic symptoms three times every 2 to 3 days by mild stress (triple stimulation technique), as described previously [37]. All *dt^sz^* hamsters used in this study exhibited severe dystonia with at least stage 3. Healthy control hamsters were treated equally. All animal experiments were carried out in accordance with the guidelines of the EU Directive 2012/63/EU and the federal laws for the protection of animals under licence Az: 7221.3-1-053/17.

### Surgical procedure for deep-brain stimulation electrode implantation and stimulation protocol

Animals (30-42 days old) were fixed in a stereotactic frame (Narishige, Japan) under deep anaesthesia with isoflurane (Isofluran, Baxter, Deerfield, IL, USA; Univentor 1200 Anaesthesia Unit + Univentor 2010 Scavenger Unit, Biomedical Instruments, Zöllnitz, Germany). The periosteum was additionally treated with the local anaesthetic bupivacaine (bupivacaine 0.25% JENAPHARM^®^). Two concentric bipolar electrodes (platinum-iridium Pt/Ir; SNEX-100, Microprobes, Gaithersburg, MD, USA) were placed bilaterally in the entopeduncular nucleus (EPN; corresponding to GPi in humans; stereotaxic coordinates AP: −0.6 mm, ML: ± 2.2 mm, DV: −0.6 mm relative to Bregma from the golden hamster atlas [44]). To provide firm fixation of the electrodes, two screws were anchored in the skull behind the electrodes and enclosed with dental adhesive (Heliobond + Compo glass flow, Schaan, Liechtenstein; SDR^®^ flow+, Dentsply DeTrey GmbH, Konstanz, Germany). After 3-5 days of recovery, the electrode wires were connected to an external programmable stimulator (Institute of Applied Microelectronics, Faculty of Computer Science and Electrical Engineering, University of Rostock) generating charge-balanced rectangular current pulses. DBS (130 Hz, 50 μA, 60 μs pulse duration) was performed for three hours on awake and freely moving animals. These parameters were chosen with regard to the proven antidystonic efficacy [43]. Every second *dt^sz^* or control hamster was used for sham stimulation (i.e. electrode implantation active stimulation) to be able to compare effects with and without stimulation. These sham-stimulated groups received the same treatment as the stimulated groups, but with the stimulator turned off.

### Brain slice preparation for analysis of striatal network excitability and inhibitory tone

Immediately after bilateral DBS or sham stimulation, the animals were decapitated under deep anaesthesia. The electrodes were carefully removed from the skull, without causing shearing movements, and the brain was quickly removed and chilled in ice-cold sucrose solution, containing (in mM): NaCl 87, NaHCO_3_ 25, KCl 2.5, NaH_2_PO_4_ 1.25, CaCl_2_ 0.5, MgCl_2_ 7, glucose 10 and sucrose 75. The brain was then cut dorsally at an angle of 40° to the horizontal axis and glued with the cut off surface to the microtome table (VT1200S, Leica Biosystems Nussloch, Germany) (Fig. 1A). The angled brain was cut horizontally in slices of 400 μm or 300 μm (field or patch clamp recordings, respectively), maintaining synaptic connections between motor cortex and striatum. After cutting, the slices (total of n=74 from control and of n=165 from dt^sz^-hamsters) were incubated for 60 min in sucrose solution at room temperature, before transferral to an interface-type recording chamber (BSC-BU, Harvard Apparatus Inc, March-Hugstetten, USA) perfused with artificial cerebrospinal fluid (ACSF) containing (in mM): NaCl 124, NaHCO_3_ 26, KCl 3, NaH_2_PO_4_ 1.25, CaCl_2_ 2.5 and glucose 10, kept constant at 32°C (TC-10, npi electronic GmbH, Tamm, Germany).

**Figure 1.**
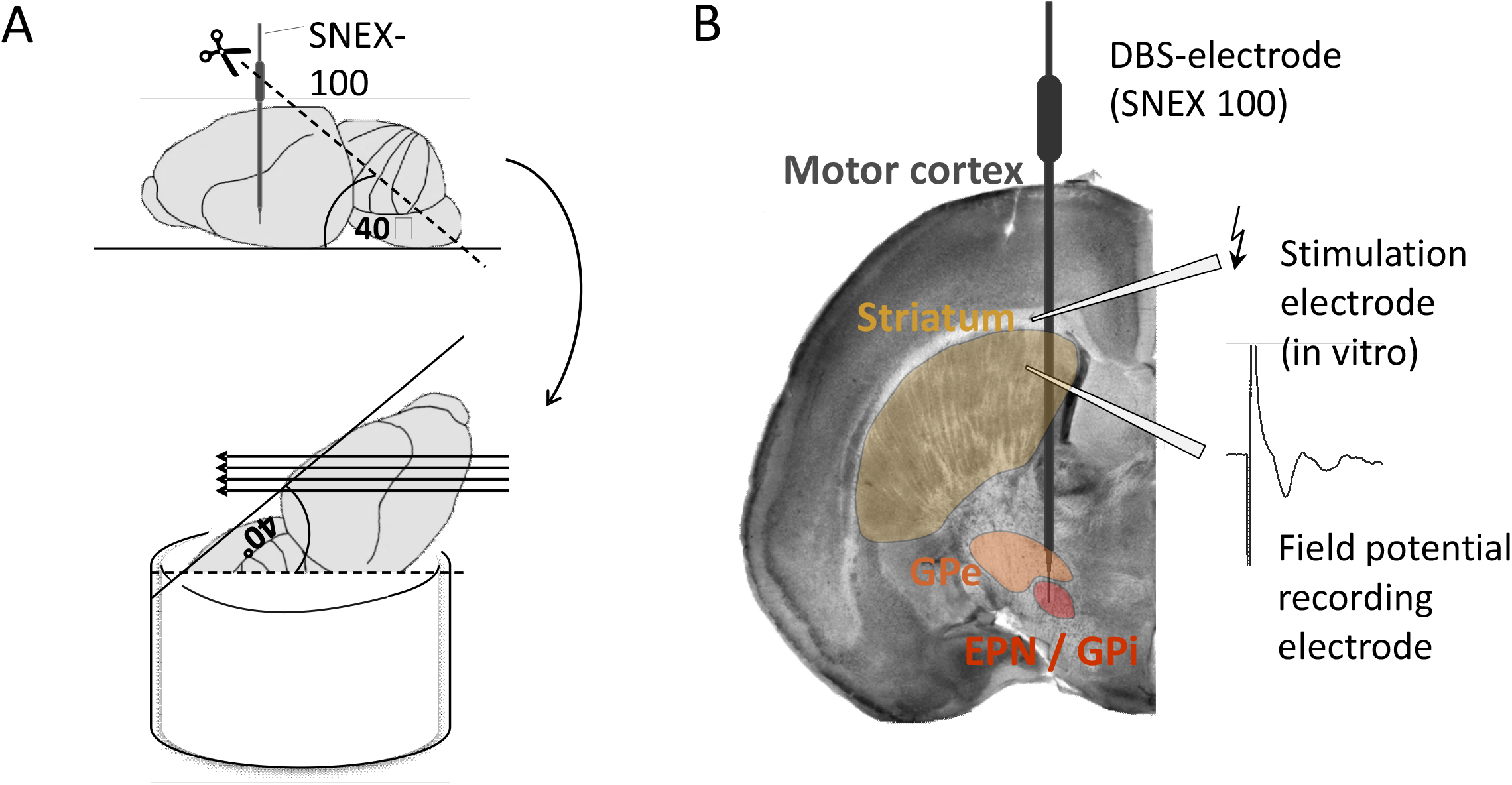
Slice generation and electrode placement. A: Illustration of brain preparation after removal from skull. Top: The position of the SNEX-100 DBS-electrode is given schematically; before slicing; the electrode was removed gently. To obtain angulated slices, and thus to preserve cortico-striatal projections, the dorsal part of the cortex and cerebellum were removed at the 40° angle as indicated by the scissors pictogram. Bottom: After angulation, horizontal cuts of the brain were performed to obtain 400 μm slices as indicated by parallel arrows. B: Photograph of a brain slice, prepared as described in the slice preparation section, containing the motor cortex and striatum, with connections between these two regions still intact. For *in vitro* recordings, the field potential recording electrode was placed in the dorso-medial part of the striatum, and the extracellular stimulating electrode (lightning arrow) was placed in the white matter of the adjacent neocortex, as indicated. Again, the virtual position of the SNEX-100 DBS electrode (now explanted) is indicated on the slice.

### Field potential recordings

Field excitatory postsynaptic potentials (fEPSP) were recorded from the dorso-medial part of the striatum using an ACSF-filled glass pipette with a silver/silver chloride wire as described before [39,45]. To activate cortical projections to the striatal network, a bipolar stimulation electrode (PT- 2T, Science Products GmbH, Hofheim, Germany) was placed into the underlying white matter of the adjacent cortex (Fig. 1B). Current-controlled stimulation was delivered by a Master-8 pulse stimulator (AMPI, Jerusalem, Israel) connected to a stimulus isolator (A365, WPI Inc, Sarasota, USA).To gauge cortico-striatal excitability, evoked input-output responses were characterised by increasing stimulus intensity stepwise until reaching saturating responses (remaining always below maximal intensity of 400 μA). For further investigations, stimulus strength was set to 50% of saturating response intensity to maintain dynamic range of responses. To test for synaptic facilitation and/or depression, paired-pulses were delivered at 40 ms inter-pulse-intervals (IPI) every 30 s. The paired-pulse ratio (PPR) was calculated by dividing the slope of the second response by the slope of the first response (s-fEPSP2/s-fEPSP1). In addition, we calculated the coefficient of variation of these synaptically evoked responses form the first 40 responses under baseline conditions in each group to be able to compare response variability. To determine the degree of inhibitory tone controlling PPR, the GABA_A_ receptor antagonist gabazine (SR 95531 hydrobromide, Tocris, Wiesbaden-Nordenstadt, Germany, 5 μM) was bath-applied for 60 min after having recorded a minimum of 20 min stable baseline responses. Signals were recorded using EXT-10-2F field-potential amplifier in AC mode (low pass filter at 1 kHz, npi, Tamm, Germany). Signals were processed and digitised at 10 kHz with Power1401 A/D converter (Cambridge Electronic Design, Cambridge, UK).

### Patch-clamp recordings

Patch-clamp recordings from medium spiny striatal neurons were obtained to assess frequency and kinetic properties of spontaneous miniature excitatory postsynaptic currents (mEPSCs) as a measure of presynaptic cortico-striatal functional modulations, and to gauge neuronal properties of medium spiny striatal neurons, identified by their characteristic firing patterns (cf. Fig. 4). Patch-clamp recordings were performed at room temperature in corticostriatal slices submerged in recording ACSF with borosilicate pipettes (3.1-8.5 MΩ, mean 5.5 ± 0.1 MΩ, n = 56, pulled with DMZ Zeitz puller, Zeitz-Instrumente Vertriebs GmbH, Martinsried, Germany) filled with a solution containing (in mM): K-gluconate 115, KCl 20, MgCl_2_ 2, HEPES 10, Na_2_-ATP 2, Mg-ATP 2, Na_2_-GTP 0.3; pH set to 7.3 and osmolarity to 280 ± 5 mosmol/l. MSN were visualized via differential interference contrast microscopy and a CCD camera (Till Photonics, Gräfelfing, Germany) enabling visual differentiation between MSN and other striatal neurons by cell shape and size. Visual classification was further re-checked by electrophysiological characterization of MSN showing specific passive and active membrane properties. The MSN recordings seals were > 1 GΩ (6.9 ± 1.8 GΩ, n = 56) and liquid junction potentials and series resistance (17.6 ± 0.7 MΩ, n = 56) were not compensated. Voltage- and current-clamp data were recorded with an EPC-10 amplifier (HEKA, Lambrecht, Germany), filtered at 1 kHz, digitized at 20 kHz and stored via Patchmaster v2.20 software (HEKA, Lambrecht, Germany). The resting membrane potential was measured initially after establishing whole cells configuration. The number of action potentials, the threshold current (rheobase) and the latency of the first action potential at rheobase were achieved at 0 pA holding current by depolarizing current injections of 500 ms duration from 0 to at least 300 pA (50 pA increments). The hyperpolarisation-activated, cyclic-nucleotide-modulated non selective (HCN) channel-dependent voltage sag was measured during hyperpolarization from a holding potential of −70 mV by current injections of 1 s duration from 0 mV to −300 pA (50 pA increments). The voltage sag amplitude was calculated as difference between the maximal hyperpolarization at the beginning and the steady state voltage at the end of the current injection. Cellular input resistance was calculated from the slope of the steady state current–voltage relation resulting from voltage steps (2 mV increments, 1 s duration) of −60 mV to −80 mV at from a holding potential −70 mV. For measurements of miniature excitatory postsynaptic currents (mEPSC) the membrane potential was clamped at −70 mV and TTX (1 μM) and gabazine (5 μM) were added to the ACSF. mEPSC events were low-pass filtered at 1 kHz and detected within 5 min with a signal to noise ratio of 5:1 using the software MiniAnalysis v.6.0.7 (Synaptosoft, Decatur, USA). Off-line analysis of patch-clamp data was performed using Fitmaster v2.11 software (HEKA), Office Excel 2003 (Microsoft, Redmond, USA) und SigmaPlot 10.0 (Systat Software GmbH, Erkrath, Germany).

### Data analysis

Data of extracellular recordings were analysed using Signal 2.16 software (Cambridge Electronic Design, Cambridge, UK). All values are given as means ± SEM; n refers to numbers of slices unless otherwise stated. Statistical analysis was performed with SigmaStat and SigmaPlot software (Systat Software Inc., San Jose, CA, USA). The significance of difference between the median values of the input-output activity of stimulated and sham-stimulated *dt^sz^* mutant and control groups were evaluated using a two-way repeated measures analysis of variance (ANOVA, two factor repetition) and a post-hoc multiple comparison procedure (Holm-Sidak method). For all other analyses, statistical significance was tested using the Wilcoxon Rank Sum Test for paired data and the Wilcoxon-Mann-Whitney Rank Sum Test for unpaired data. A probability value of P < 0.05 was considered significant indicated by asterisks (* unpaired test) and by hash (# paired test) respectively.

## Results

The aim of this study was to explore the possible mechanisms underlying the antidystonic effect of 3h-DBS delivered to freely moving animals reported recently by our group [43]. Our focus was on exploring changes in cortico-striatal communication, since dystonias are thought to involve a disturbance in the balance of striatal control of the GPi [1,6], the equivalent of the entopeduncular nucleus (EPN) in rodents.

### Input-output relationship of cortico-striatal synaptic connections

We were first interested whether EPN-DBS changed the overall efficacy of synaptic connectivity between motor cortex and striatum. To gauge this, we explored the so-called input-output relationship of evoked field potential responses in the dorso-medial striatum to cortical activation via local afferent fibre stimulation. For this, stimulation intensity was stepwise increased from threshold to saturating response. As Fig. 2A illustrates, the resulting field excitatory postsynaptic potentials (fEPSP) increased in amplitude with cumulative rising stimulus in both groups, healthy (WT; triangles) and dt^sz^ (circles). Of note, without DBS (empty symbols), the magnitudes of the responses in healthy tissue were indeed comparable to, but somewhat smaller than in dt^sz^, with maxima at around 0.4 V/s (control) to 0.5 V/s (dt^sz^), as already shown previously [46]. As Fig. 2A shows, EPN-DBS (filled symbols) altered the responsiveness of this cortico-striatal synapse, but only in healthy tissue, where it essentially doubled the slope of the responses (p<0.05, ANOVA). Specifically, the mean values of field potential slopes (in V/s) were −0.56 ± 0.06 (*dt^sz^*, n=65) and −0.44 ± 0.06 (control, n=14) for sham-stimulated tissue, and −0.61 ± 0.08 V (*dt^sz^*, n=42) and −0.96 ± 0.20 V (control, n=19) for EPN-DBS-stimulated groups, as a response to the highest cortico-striatal stimulus intensity of 400 μA. Thus, DBS enhances synaptic efficiency only in healthy, wild-type tissue.

**Figure 2.**
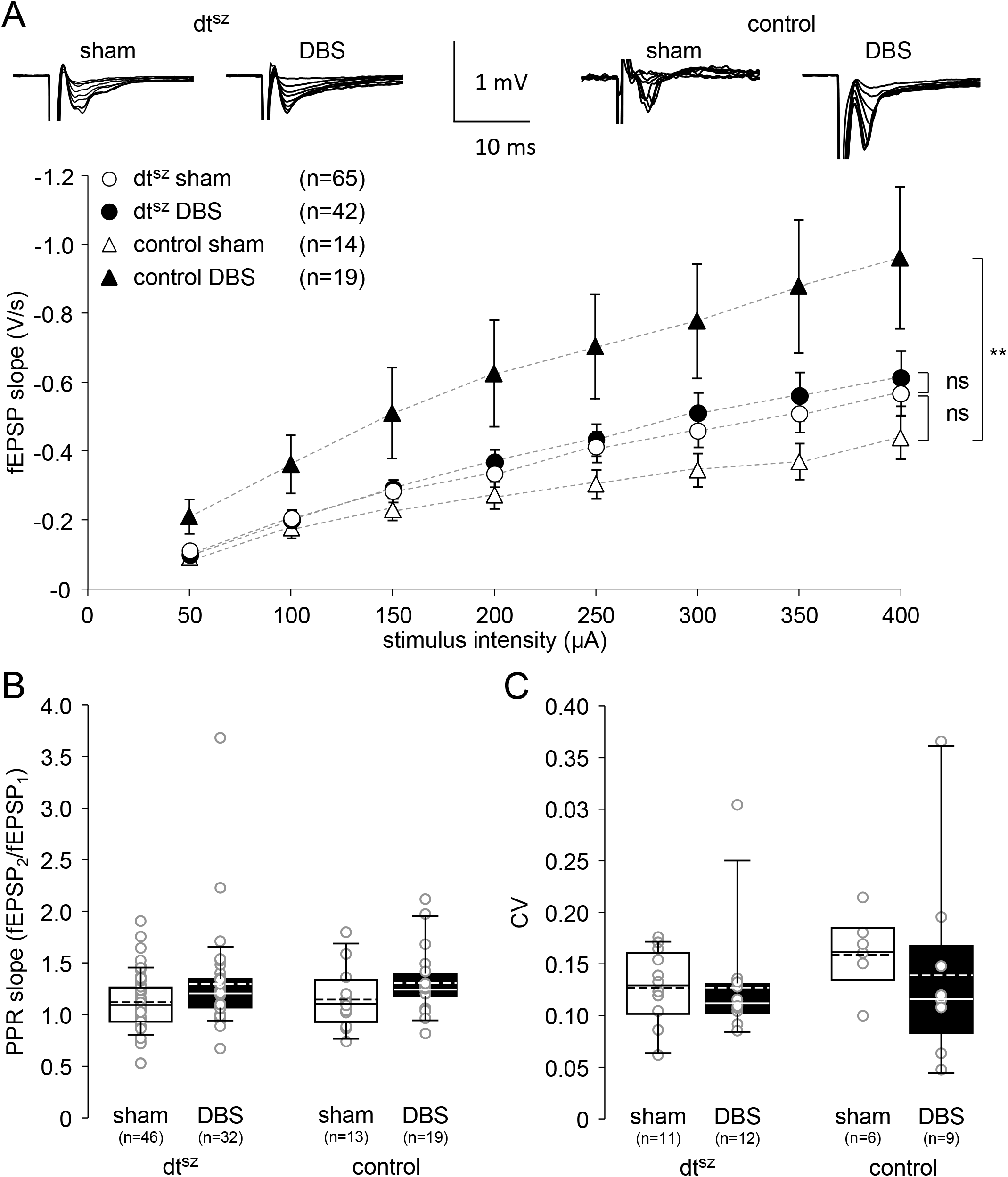
DBS enhances synaptic efficiency in control, but not in the dystonic groups. **A:** The dot plot shows the effect of DBS on input-output behaviour in the cortico-striatal network. Average data of pooled fEPSP slopes in response to increasing stimulating of the four experimental groups are indicated by dots and slashed lines. Data are presented as mean ± SEM of stimulated (filled symbols) and sham-stimulated (empty symbols) control (triangles) and *dt^sz^* hamsters (circles). The number of slices is given in parentheses (in each experiment, usually only one slice per animal was obtained). Asterisks indicate significant differences (p < 0.01; ANOVA). Representative traces (top) illustrate series of fEPSP in response to increasing stimulus strengths of the respective groups as indicated. **B:** Box and whisker plot of paired-pulse ratio (PPR) of evoked field potential slopes in dtsz and control slices, without DBS (sham), and after EPN-DBS (DBS). A PPR > 1 demonstrates facilitation, and <1 depression of the second of a pair of responses evoked at an interval of 40 ms. Medians: straight lines, means: hashed lines. Single dots represent means of one experiment (slice). **C**: Box and whisker plot of coefficient of variation of evoked field potential slopes in dtsz and control slices, without DBS (sham), and after EPN-DBS (DBS). Means: straight lines, medians: hashed lines. Single dots represent means of one experiment (slice).

### Variability of cortico-striatal synaptic responses

Since DBS is speculated to normalise bursting oscillatory activity [47] and to disrupt aberrant synaptic transmission [48], we hypothesised that cortico-striatal responses would show greater variability in dystonic tissue, which should be reduced by DBS. As shown in Fig. 2C, the coefficients of variation of the synaptic responses (all in the range of 0.13 to 0.16, see Table 1 for details) did not differ between control and dystonic tissue, and remained unaffected by EPN-DBS. Hence, we had to dismiss both hypotheses.

**Table 1.** Variability of evoked compound synaptic cortico-striatal potentials: Coefficient of variation.

3 Variability of evoked compound synaptic cortico-striatal potentials: Coefficient of variation
| **dt<sup>sz</sup>** |  | **control** |  |
| --- | --- | --- | --- |
| **sham** | **DBS** | **sham** | **DBS** |
| **12.7 ± 1.0**<br>(n=11) | **12.7 ± 1.6**<br>(n= 12) | **15.9 ± 1.4**<br>n=6 | **13.9 ± 3.0**<br>n=9 |

4 Values are means ± SEM

### Paired-pulse ratio (PPR)

We were also interested in the effect of EPN-DBS on short-term plasticity at the cortico-striatal synapse, since this plasticity governs the fidelity of transmission of repetitive synaptic events. For this, we gauged paired-pulse responses at the cortico-striatal synapse using fEPSP as before, now elicited in short succession twice at 40 ms inter-stimulus intervals. Paired pulse response changes (facilitation or depression of the second response) are generally thought to be based on presynaptic release probability alterations, with paired-pulse depression (PPD) probably reflecting a presynaptic Ca^2+^-dependent effect on release probability which however is under modulatory tone of GABA and can thus be reduced with loss of inhibition [49]. Paired-pulse facilitation (PPF), in turn, is supposed to be caused by an initially low release probability, which increases with residual presynaptic Ca^2+^ [50,51]. PPF was present in all groups (Fig. 2B). Thus, the ratio 2^nd^/1^st^ pulse was 1.15 ± 0.08 (control tissue, sham DBS; n=13), 1.31 ± 0.07 (control tissue, DBS; n=19), 1.12 ± 0.04 (*dt^sz^* tissue, sham DBS; n=46) and 1.30 ± 0.09 (*dt^sz^* tissue, DBS; n=32); these values did not differ significantly, even though the values under DBS were always higher than those without – an effect of DBS thus cannot be ruled out completely.

### Inhibitory control of cortico-striatal synaptic communication

With regard to evidence of reduced GABAergic inhibition of striatal projection neurons, probably based on deficient striatal GABAergic interneurons in dystonic hamsters [33–35] and considering the postulated GABAergic disinhibition in patients with dystonia, where a shift of balance toward the indirect pathway is speculated to occur [6], we were interested whether the synaptic responses evoked in the striatum by cortical afferents would be under the control of GABAergic inhibition, and whether this GABAergic control might change after DBS. We therefore explored the reaction of evoked fEPSP under the blockade of GABA_A_ receptors using gabazine (5 μM) application. As illustrated in Fig. 3A (dot plot of fEPSP responses during continuing gabazine application) and Fig. 3B (example traces of fEPSP before and after GABA-block), suppressing GABA_A_-receptor activation indeed had the effect of increasing the striatal field responses by 2-3 fold. Notably, this was more significant in control than in dystonic, *dt^sz^* tissue (to 309.8 ± 34.3 % vs. 250.4 ± 34.5 % of control values, respectively, p<0.05, means ± SEM of values of last 5 min, controls n=6, dt^sz^ n=11), suggesting inhibitory control of synaptic activity to be higher in non-dystonic controls than in dystonic animals. Importantly, this inhibitory control as evidenced by the gabazine effect was significantly modulated by DBS, and differentially so in control vs. dystonic group: While in non-dystonic controls, DBS was associated with a significantly lower increase in fEPSP slope compared to sham-DBS stimulation (p<0.05, ANOVA), the contrary was the case in dystonic, *dt^sz^* tissue, where DBS led to a higher increase in fEPSP (p<0.05, ANOVA). Thus, in the control group, the slope fell from 309.8 ± 34.3 to 263.6 ± 56.1. In slices from *dt^sz^*-hamsters, by contrast, the slope rose from 250.4 ± 34.5 to 282.3 ± 44.0 (controls DBS n=9, dt^sz^ DBS n=12, means ± SEM of values of last 5 minutes).

**Figure 3.**
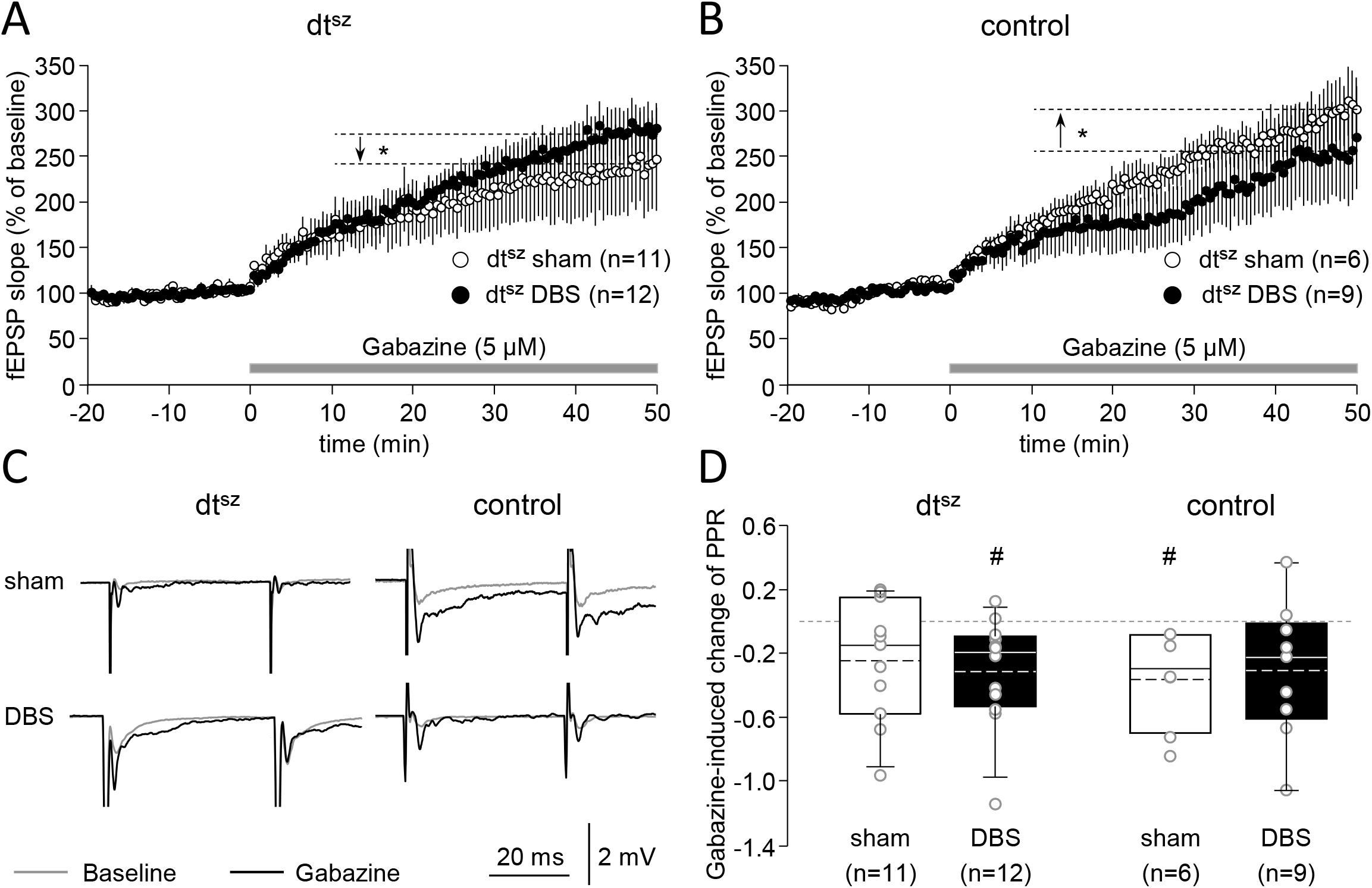
DBS specifically increases striatal inhibitory tone in dystonic animals, and decreases it in non-dystonic hamsters. **A, B:** The dot plots show the effect of DBS on cortico-striatal synaptic transmission under GABA-receptor blockade using gabazine (5 μM) from time-point 0, as indicated, after 20 min of stable baseline conditions. The data points represent cumulative means of relative fEPSP changes as percentage of baseline values before GABA-receptor block. Relative increases of fEPSP slope under GABA-receptor block hence indicate degree of inhibitory tone controlling synaptic transmission. fEPSP slopes were measured at 50% saturating stimulus intensity. Data are presented as mean ± SEM of EPN-DBS stimulated (filled symbols) and sham-stimulated (empty symbols) in dt^sz^ (**A**) and control hamsters (**B**). The number of slices is given in parentheses. Asterisks indicate significant differences (p < 0.05; two-way repeated measurement ANOVA) and refer to comparisons of EPN-DBS stimulated (DBS) and sham-stimulated (sham) animals. **C**: Representative traces illustrate fEPSP in slices of dt^sz^ and control animals within the first (fine trace, baseline, t = −20 to −15 min) and last five min (bold trace, gabazine, t= 55 to 60 min) of the measurement. **D**: Box and whisker plot of changes of paired-pulse ratio (PPR) after GABA_A_-receptor block (gabazine) coefficient of variation of evoked field potential slopes in dt^sz^ and control slices, without DBS (sham), and after EPN-DBS (DBS). In all plots, filled symbols represent data from animals having undergone EPN-DBS, open symbols those of animals with sham stimulation only. In all box plots, medians are represented by straight lines and means by dashed lines. Single dots represent means of one experiment (slice).

We were also interested in which way the PPR would be modulated by GABA_A_-receptor inhibition (5 μM gabazine, 20 min application). In all groups, the PPR was reduced during GABA_A_ block, i.e. the second of these paired responses became similar to the first, or even smaller than it. This suggests that GABAergic tone apparently also dampens presynaptic release probability at this synapse. Thus, the reduction amounted to 0.36 ± 0.11 (control tissue, sham DBS; n=11), 0.31 ± 0.13 (control tissue, DBS; n=9), 0.24 ± 0.11 (*dt^sz^* tissue, sham DBS; n=11) and 0.31 ± 0.09 (*dt^sz^* tissue, DBS; n=12). Interestingly, the reduction was significant (p<0.05, MWRS-test) for healthy tissue only under control conditions without DBS, and for dystonic tissue only after DBS. Thus, facilitation of responses reversed to depression in these cases, again supporting the notion that inhibitory control in healthy tissue is reduced after DBS in healthy tissue, and increased after DBS in dystonic one.

### Spontaneous cortico-striatal synaptic activity

The field potential investigations so far remained on a compound network level. We therefore strove to look at glutamatergic cortico-striatal synapses in more detail, i.e. on the single-cell level, by analysing miniature excitatory postsynaptic currents (mEPSC), reflecting spontaneous release activity from cortical projections. As shown in Fig. 4, EPN-DBS had a strong effect on the frequency of mEPSC, reducing it in both healthy and dystonic animals. Interestingly, this reduction was stronger in dystonic tissue, and indeed significant only in this case. Thus, the frequency of mEPSC after EPN-DBS fell from 4.11 ± 0.39 Hz to 2.15 ± 0.69 Hz (p<0.05, MWRS-test) in dystonic tissue, and from 3.18 ± 0.5 to 1.70 ± 0.45 in control (n.s.) (Fig. 4A and C). At the same time, neither amplitudes, nor rise or decay times of the mEPSC (Fig. 4 B2-4 and D2-4) differed among the groups, even though the peak incidence in dystonic tissue shifted from 6 to 8 pA after EPN-DBS (Fig. 4D1), while it remained at 7 pA in healthy tissue (Fig. 4B2) (for details on the values, see Table 2). EPN-DBS thus obviously dampens spontaneous presynaptic glutamate release at cortico-striatal synapses, and this again differentially stronger in dystonic than in healthy tissue.

**Figure 4.**
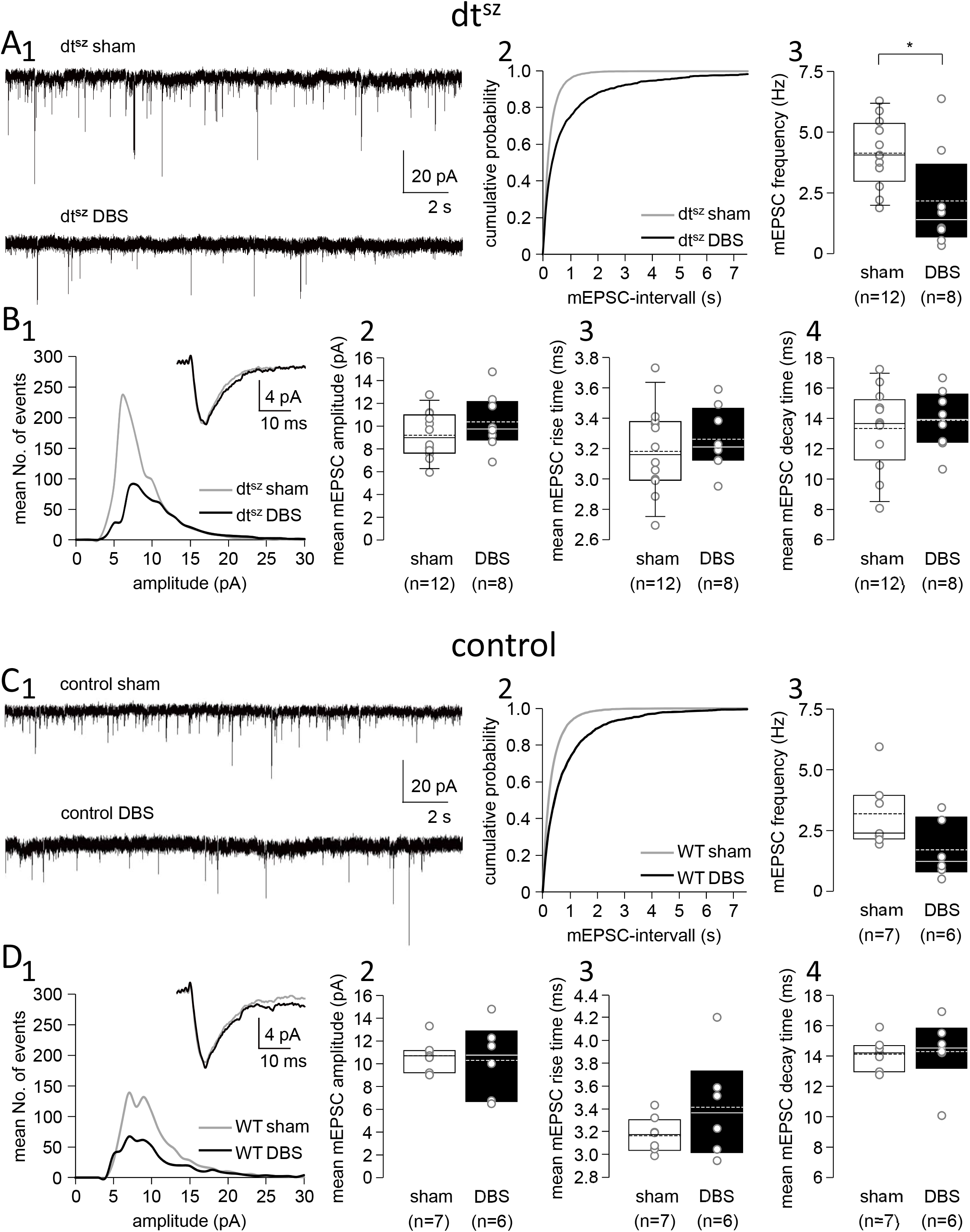
DBS decreases miniature EPSC (mEPSC) occurring spontaneously at the cortico-striatal synapse of medium spiny neurones. Original traces (**A1, C1**) of mEPSCs recorded from medium spiny neurones in dt^sz^ (**A**) and control (**C**) slices from animals having undergone EPN-DBS (DBS) or sham stimulation (sham). Corresponding cumulative probability histograms (**A2, C2**) as well as box and whisker plots (**A3, C3**) of the same groups (sham: light curves and open box plots, dt^sz^: dark curves and filled boxes) show that DBS reduces frequency of mEPSC, with a significant reduction in dystonic tissue (asterisk in A3, p<0.05, MWRS test). By contrast, amplitudes of mEPSC are not altered by DBS with respect to sham stimulation in either dystonic (dt^sz^, **B**) or control slices (control, **D**), as illustrated by amplitude distribution histograms (**B1, D1**), as well as box and whisker plots of mean mEPSC amplitudes (**B2, D2**). Neither are there any differences among the groups regarding the kinetics of mEPSC, as illustrated in the box and whisker plots of mEPSC mean rise (**A3, D3**) and decay (**A4, D4**) times. In all plots, filled symbols represent data from animals having undergone EPN-DBS, open symbols those of animals with sham stimulation only. In all box plots, medians are represented by straight lines and means by dashed lines. Single dots represent means of one experiment (slice).

**Table 2.** Characteristics of spontaneous miniature excitatory currents on medium-spiny neurons.

6 Characteristics of spontaneous miniature excitatory currents on medium-spiny neurons
|  | **dt<sup>sz</sup>** |  | **control** |  |
| --- | --- | --- | --- | --- |
|  | **Sham**<br>n=12 | **DBS**<br>n=8 | **Sham**<br>n=7 | **DBS**<br>n=6 |
| **mEPSC frequency (HZ)** | **4.11 ± 0.39** | **2.15 ± 0.69** | **3.18 ± 0.50** | **1.70 ± 0.45** |
| **mEPSC amplitude (pA)** | **9.20 ± 0.56** | **10.39 ± 0.82** | **10.68 ± 0.50** | **10.70 ± 1.22** |
| **mEPSC rise time (ms)** | **3.18 ± 0.08** | **3.26 ± 0.07** | **3.17 ± 0.05** | **3.42 ± 0.17** |
| **mEPSC decay time (ms)** | **13.33 ± 0.75** | **13.87 ± 0.66** | **14.14 ± 0.37** | **14.30 ± 0.17** |

Values are means ± SEM

### Neuronal properties

Last, we were interested in the effect of EPN-DBS on the postsynaptic level, i.e. on cellular properties of medium spiny neurons receiving cortical input. We thus tested the firing properties of these neurons upon intracellular current injection at increasing strengths (Fig. 5 A, D), the so-called voltage sag presumably mediated by hyperpolarisation-activated, cyclic-nucleotide-modulated non selective (HCN) channels (which modulate excitability) (Fig. 5 B, E), as well as input resistance, membrane capacitance, resting membrane potential, rheobase and latency to first AP (which characteristically is >50 ms in these neuron types) (box plots in Fig. 5C and F). There were no differences, neither between healthy (control) and dystonic (dt^sz^) tissue, as already reported before in this dystonia model [46], and actually also a mouse DYT1 model [52], nor between conditions without (sham) or with EPN-DBS (DBS) (for details on the values, see Table 3). Hence, we can conclude that EPN-DBS does not alter striatal medium spiny neuron properties.

**Figure 5.**
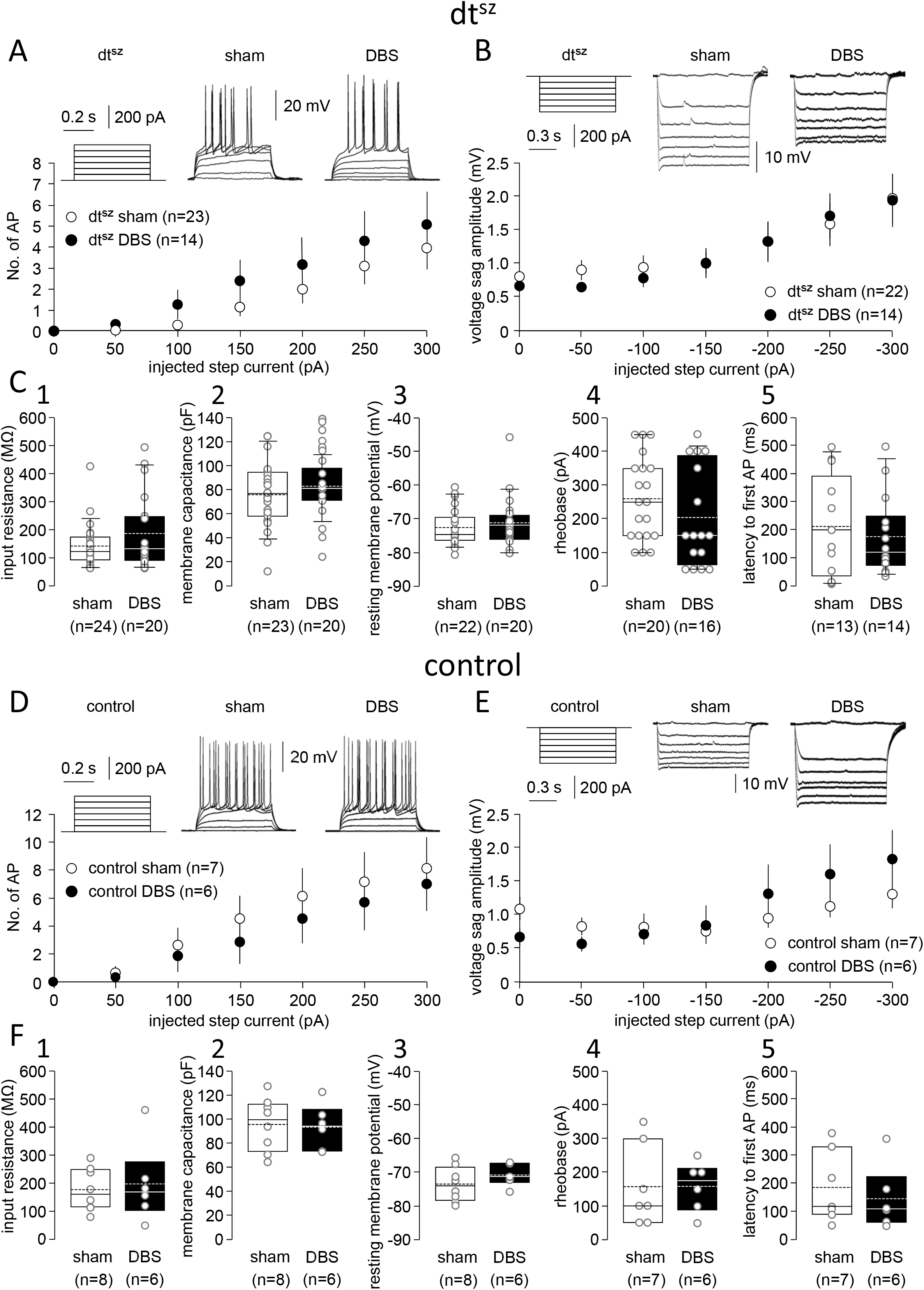
DBS does not influence intrinsic neuronal properties of medium spiny neurones. Original traces of membrane potential recordings illustrating firing properties (**A, D**) of medium spiny neurones in dt^sz^ (**A**) and control (**D**) slices from animals having undergone EPN-DBS (DBS) or sham stimulation (sham). Firing was elicited by depolarising current injection at increasing amplitudes (inset left). The corresponding input-output curve is displayed below the original traces, as dot diagram of action potential number plotted against current injection. Original traces of membrane potential recordings illustrating voltage sag (**B, E**) of medium spiny neurones in dystonic (dt^sz^, **A**) and control (**D**) slices from animals having undergone EPN-DBS (DBS) or sham stimulation (sham). Voltage sag representing hyperpolarisation-activated, cyclic-nucleotide gated, non-selective (HCN) channel activation was elicited by hyperpolarising current injection at increasing amplitudes (inset left). The corresponding input-output curve is displayed below the original traces, as dot diagram of voltage change in depolarising direction plotted against current injection. **C, F**: Box and whisker plots of input resistance (**C1, F1**), membrane capacitance (**C2, F2**), resting membrane potential (**C3, F3**), rheobase (**C4, F4**) and latency to first action potential after depolarising current injection (**C5, F5**) in tissue from dystonic (**C**) and control (**F**) animals In all plots, filled symbols represent data from animals having undergone EPN-DBS, open symbols those of animals with sham stimulation only. In all box plots, means are represented by straight lines and medians by dashed lines. Single dots represent means of one experiment (slice). In all plots, filled symbols represent data from animals having undergone EPN-DBS, open symbols those of animals with sham stimulation only. In all box plots, medians are represented by straight lines and means by dashed lines. Single dots represent means of one experiment (slice).

**Table 3.** Properties of medium spiny striatal neurones.

|  | **dt<sup>sz</sup>** |  | **control** |  |
| --- | --- | --- | --- | --- |
|  | **Sham** | **DBS** | **Sham** | **DBS** |
| **maximum no. of AP (at 300 pA)** | **4.0 ± 1.0**<br>n=10 | **5.1 ± 1.6**<br>n=13 | **8.13 ± 2.18**<br>n=8 | **7.00 ± 1.90**<br>n=6 |
| **maximum of voltage sag amplitude (at -300 pA) (mV)** | **1.96 ± 0.37**<br>n=14 | **1.93 ± 0.39**<br>n=22 | **1.31 ± 0.20**<br>n=7 | **1.83 ± 0.43**<br>n=6 |
| **input resistance (MΩ)** | **142.5 ± 15.6**<br>n=24 | **185.5 ± 28.9**<br>n=20 | **177.2 ± 25.2**<br>n=8 | **197.5 ± 52.5**<br>n=6 |
| **membrane capacitance (pF)** | **76.1 ± 5.6**<br>n=23 | **82.9 ± 4.2**<br>n=20 | **95.67 ± 7.33**<br>n=8 | **95.52 ± 7.05**<br>n=6 |
| **resting membrane potential (mV)** | **-72.6 ± 1.1**<br>n=22 | **-71.1 ± 1.7**<br>n=20 | **-73.5 ± 1.7**<br>n=8 | **-70.8 ± 1.21**<br>n=6 |
| **rheobase (pA)** | **260.0 ± 26.8**<br>n=20 | **203.1 ± 35.8**<br>n=16 | **157.1 ± 42.2**<br>n=8 | **158.3 ± 27.4**<br>n=6 |
| **Latency of 1<sup>st</sup> AP (ms)** | **212.2 ± 47.0**<br>n=13 | **176.8 ± 36.6**<br>n=14 | **185.3 ± 44.6**<br>n=8 | **145.1 ± 42.6**<br>n=6 |

Values are means ± SEM

## Discussion

In this study we could show that DBS of the EPN for 3h in awake animals, which leads to alleviation of dystonic symptoms in mutant hamsters, which we could demonstrate recently [43], is associated with functional changes in cortico-striatal synaptic communication.

### Properties defining the dystonic condition

Studies in patients suggest that a pathological cortico-striatal function, and subsequent disturbance of striatal control of GPi, are likely to be important factors of dystonic dysfunction [1], possibly resulting in a shift of the balance toward the direct pathway [6]. In this sense, the animal models of dystonia, and in particular the dystonic *dt^sz^* mutant hamster, mirror this condition: It is spontaneously dystonic, and, as speculated for at least some human dystonias [3,4] [5], displays increased cortico-striatal excitability, as a result of reduced intra-striatal GABAergic signalling [35],[39]. This is corroborated by findings in humans, where reduced cortical and striatal inhibition were reported [53]. Even though there is an apparent in contrast to monogenetic dystonias such as DYT1 models, where GABAergic transmission was actually found to be reduced [52], the fact that instead inhibition via cholinergic interneurones was reverted to excitation in these models [54] also substantiates a deficit in inhibition, albeit via a different cell type. One can thus hypothesise that the dystonic phenotype arises from a functional shift in basal ganglia circuitry, which originates from a disinhibited striate body, which in turn results in a more prominent pallidal / entopeduncular inhibition as schematically shown in Fig. 6A.

**Figure 6.**
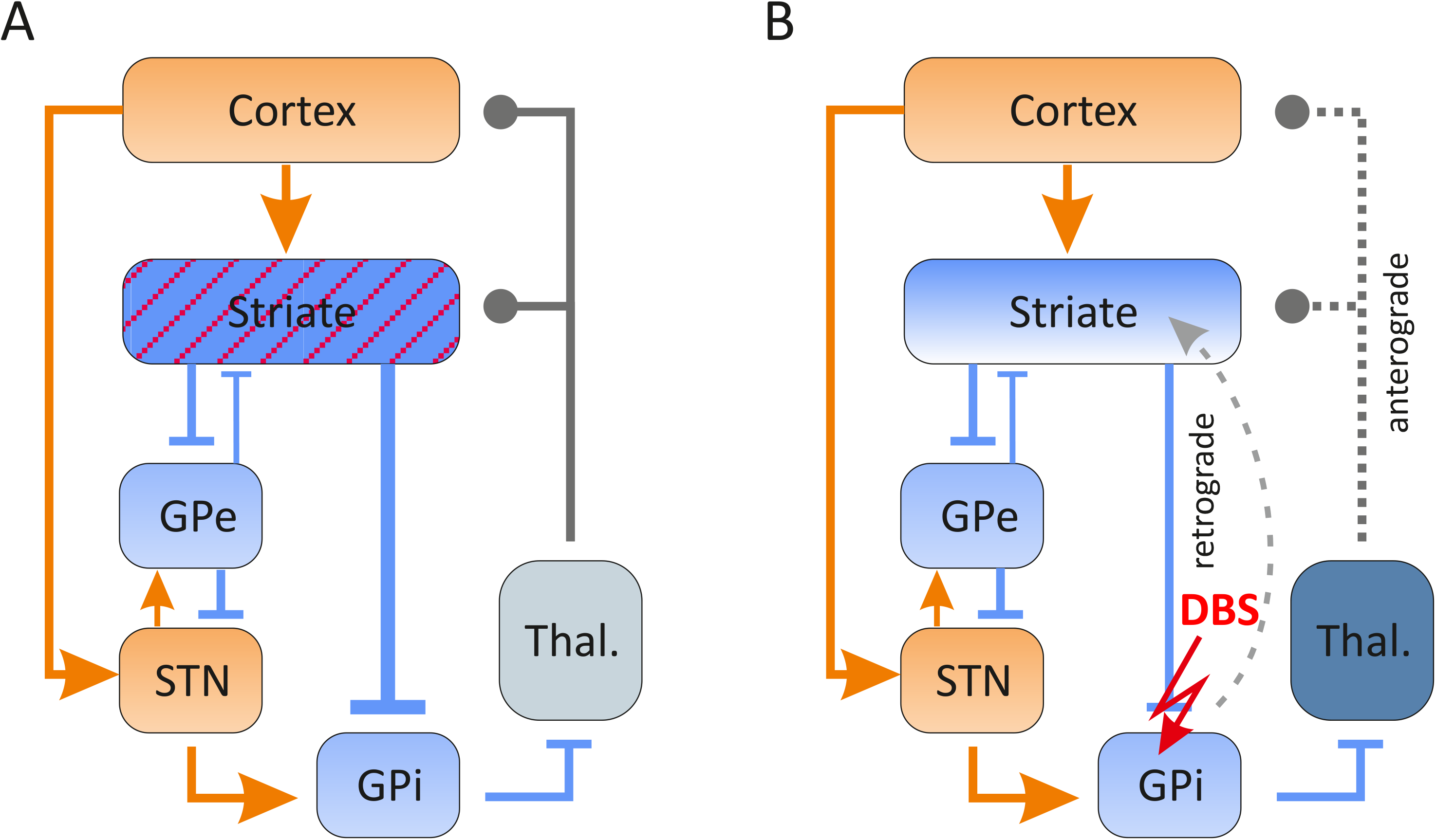
Hypothetical effect of DBS on basal ganglia circuitry. The graph shows a schematic and reduced representation of basal ganglia circuitry under dystonic conditions before (A) and after DBS (B). A: Previous findings in animal dystonia [35,42]models and data from human studies [13]suggest a loss of inhibitory tone within the striatal network, possibly based on a reduction of interneurone function, which at least in the *dt^sz^* hamster has been documented by transient loss of parvalbumin-positive interneurones [35]. We hypothesise that this leads to a disinhibition within the striatum (red hatching), and hence a more prominent inhibitory projection onto the GPi (bold lines in blue). B: We hypothesise that DBS (red lightning arrow) normalises this state, possibly via activating backfiring axons into the Striate (dotted light grey arrow), or more likely via anterograde signalling via thalamic projections onto striatum or pallido-thalamo-cortical loop (dark grey arrows).

### DBS alters cortico-striatal communication: synaptic efficacy

Our experiments demonstrate that EPN-DBS increases cortico-striatal evoked compound synaptic potentials, but only from healthy, control animals. How does DBS then change this circuitry? From patient studies, comparatively few data exist on activity changes brought about by pallidal DBS within the basal ganglia network. Pallidal stimulation suppresses low-frequency activity in the pallidum itself, and this low-frequency activity is a persistent marker of disease severity [55]. In addition, pallidal neurones react either with a persistent increase of activity, or with a sequence of events comprising initial increase, and prolonged decrease of spiking [56]. From a case study on one patient, we know that pallidal DBS in turn generates complex downstream effects on thalamic neuronal firing, with close to 48% of neurones showing a decrease in discharge frequency, and the rest an increase (8%) or no change (44%) [26]. Further downstream, pallidal DBS also seems to increase motor cortical inhibition [25]. Since there is both a thalamic, cortical and indeed pallidal, retrograde axonal connection to the striatum, effects on cortico-striatal communication have been speculated on [57], but have not been reported so far. Also animal studies do not directly address the question: Pallidal spiking changes have been confirmed in healthy rodent tissue in vitro, with the biphasic responses being attributed to cholinergic modulation [56]. Again, animal studies directly investigating cortico-striatal communication are lacking. We are hence left to speculate that the enhancing effect of EPN-DBS exclusively in healthy animals is related to the apparently differential effects on GABAergic tone (see below).

### DBS alters striatal inhibitory tone

In our study, we could show that EPN-DBS differentially affects inhibitory tone in healthy (relative reduction in tone) and dystonic tissue (relative increase in tone). This differential effect certainly is highly interesting, and could constitute one important factor in the mechanism of DBS. It is tempting to speculate that this inhibitory tone modulation results in a normalisation of the intrastriatal inhibition (which in dystonic tissue was shown to be abnormal) (Fig. 6B), although obviously we cannot rule out that also changes in feed-forward inhibition coming from the cortex contributes to this effect. Specifically, the observation that EPN-activity is reduced in *dt^sz^* hamsters [35] is very likely due to striatal overactivity in dystonic animals, and previous findings showing that intrastriatal injection of GABA blockers worsen dystonia [58] also stress the pivotal role of intrastriatal GABAergic control. We therefore speculate that the inherent loss of parvalbumin-positive interneurones is functionally alleviated by DBS, but alternatively, also a possible pathological contribution of cholinergic interneurons being overactive and hence activating medium-spiny neurones [59] might be normalised. At any rate, the paired-pulse experiments support this notion: facilitation reverted to depression in control sham, and DBS dystonic tissue under GABA block: Again, this corroborates that GABAergic tone controls synaptic transmission, and that EPN-DBS seems to reinstate a stronger GABAergic control on synaptic transmission in dystonic tissue, and that by contrast, DBS reduces this GABAergic containment in non-dystonic tissue. A caveat is that all these measurements are somewhat indirect – studies in the future thus have to address will have to disclose whether this effect is exerted directly, as known for dopaminergic synapses [60], or indirectly, by dissecting the different roles of local GABAergic and cholinergic interneurones, and feed-forward cortical inhibition.

### DBS alters spontaneous release in cortico-striatal synapses on medium spiny neurones

A prominent effect of DBS in the current study is the decrease of mEPSC frequency in tissue from dystonic animals, and to a lesser degree from healthy animals having undergone EPN-DBS. From studies in dystonia, no reports are available on this phenomenon. However, the reaction bears similarities to an effect of very high frequency spinal cord stimulation, which reduces mEPSC frequency in lamina II dorsal horn neurones to normal values [61]. How is this effect mediated? Regarding our data on intrinsic properties of the postsynaptic, medium spiny neurones, which remained unaltered by EPN-DBS, a retrograde effect of the axons projecting to the EPN backfiring into the striate (as indicated by the arrow in Fig. 6B) is unlikely, although it cannot be fully excluded. A more plausible hypothesis would be that either direct thalamic projections to the striatum or indeed indirect projections via the pallido-thalamo-cortical loop are responsible. Again, these issues await exploration in the future by exploring thalamic and cortical changes.

## Acknowledgement

We are grateful for the excellent technical assistance and animal keeping respectively of Tina Sellmann, Imke Reich, Hanka Schmidt and Silke Birkmann.

## Author contributions

RK and AR designed the study, FP, CN and DT designed and constructed the DBS stimulator, MP,DF,FP,CN,DT,CB and UvR proposed the layout of the stimulator features, FP,CN,DT as well as MZ, DF, VN, MP, SP, AL, AR and RK tested it in pilot conditions, MH, MZ, and DF conducted in vivo DBS for this study, and conducted the electrophysiological experiments, MZ,MH,DF and RK wrote the paper and all authors contributed to discussion of the manuscript.

## Highlights

- Pallidal DBS was applied in an animal model of dystonia in freely moving animals for 3h
- Persistent effects on cortico-striatal synaptic communication were observed
- DBS increased striatal inhibitory tone in dystonic, and decreased it in non-dystonic tissue.
- DBS further leads to reduction of spontaneous excitatory cortico-striatal activity in dystonic tissue.
- We hypothesise that these DBS effects are probably mediated by presynaptic modulation of cortical afferents.

## Reference List

[1] Berardelli A, Rothwell JC, Hallett M, Thompson PD, Manfredi M et al. The pathophysiology of primary dystonia. Brain 1998; 121 (Pt 7):1195–1212.

[2] Marsden CD, Harrison MJ, Bundey S. Natural history of idiopathic torsion dystonia. Adv Neurol 1976; 14:177–187.

[3] Quartarone A, Hallett M. Emerging concepts in the physiological basis of dystonia. Mov Disord 2013; 28:958–967. 10.1002/mds.25532 [doi].

[4] Quartarone A, Pisani A. Abnormal plasticity in dystonia: Disruption of synaptic homeostasis. Neurobiol Dis 2011; 42:162–170.

[5] Schirinzi T, Sciamanna G, Mercuri NB, Pisani A. Dystonia as a network disorder: a concept in evolution. Curr Opin Neurol 2018; 31:498–503. 10.1097/WCO.0000000000000580 [doi].

[6] Wichmann T, Dostrovsky JO. Pathological basal ganglia activity in movement disorders. Neuroscience 2011; 198:232–244. S0306-4522(11)00732-9 [pii];10.1016/j.neuroscience.2011.06.048 [doi].

[7] Meunier S, Russmann H, Shamim E, Lamy JC, Hallett M. Plasticity of cortical inhibition in dystonia is impaired after motor learning and paired-associative stimulation. Eur J Neurosci 2012; 35:975–986. 10.1111/j.1460-9568.2012.08034.x [doi].

[8] Beck S, Hallett M. Surround inhibition in the motor system. Exp Brain Res 2011; 210:165–172. 10.1007/s00221-011-2610-6 [doi].

[9] Beck S, Shamim EA, Richardson SP, Schubert M, Hallett M. Inter-hemispheric inhibition is impaired in mirror dystonia. Eur J Neurosci 2009; 29:1634–1640. EJN6710 [pii];10.1111/j.1460-9568.2009.06710.x [doi].

[10] Crowell AL, Ryapolova-Webb ES, Ostrem JL, Galifianakis NB, Shimamoto S et al. Oscillations in sensorimotor cortex in movement disorders: an electrocorticography study. Brain 2012; 135:615–630. awr332 [pii];10.1093/brain/awr332 [doi].

[11] Kuhn AA, Brucke C, Schneider GH, Trottenberg T, Kivi A et al. Increased beta activity in dystonia patients after drug-induced dopamine deficiency. Exp Neurol 2008; 214:140–143. S0014-4886(08)00305-1 [pii];10.1016/j.expneurol.2008.07.023 [doi].

[12] Silberstein P, Kuhn AA, Kupsch A, Trottenberg T, Krauss JK et al. Patterning of globus pallidus local field potentials differs between Parkinson’s disease and dystonia. Brain 2003; 126:2597–2608. 10.1093/brain/awg267 [doi];awg267 [pii].

[13] Tisch S, Rothwell JC, Bhatia KP, Quinn N, Zrinzo L et al. Pallidal stimulation modifies after-effects of paired associative stimulation on motor cortex excitability in primary generalised dystonia. Exp Neurol 2007; 206:80–85. S0014-4886(07)00139-2 [pii];10.1016/j.expneurol.2007.03.027 [doi].

[14] Balint B, Mencacci NE, Valente EM, Pisani A, Rothwell J et al. Dystonia. Nat Rev Dis Primers 2018; 4:25. 10.1038/s41572-018-0023-6 [doi];10.1038/s41572-018-0023-6 [pii].

[15] Prudente CN, Hess EJ, Jinnah HA. Dystonia as a network disorder: what is the role of the cerebellum? Neuroscience 2014; 260:23–35. S0306-4522(13)01009-9 [pii];10.1016/j.neuroscience.2013.11.062 [doi].

[16] Krack P, Volkmann J, Tinkhauser G, Deuschl G. Deep Brain Stimulation in Movement Disorders: From Experimental Surgery to Evidence-Based Therapy. Mov Disord 2019; 34:1795–1810. 10.1002/mds.27860 [doi].

[17] Volkmann J, Mueller J, Deuschl G, Kuhn AA, Krauss JK et al. Pallidal neurostimulation in patients with medication-refractory cervical dystonia: a randomised, sham-controlled trial. Lancet Neurol 2014; 13:875–884. S1474-4422(14)70143-7 [pii];10.1016/S1474-4422(14)70143-7 [doi].

[18] Volkmann J, Wolters A, Kupsch A, Muller J, Kuhn AA et al. Pallidal deep brain stimulation in patients with primary generalised or segmental dystonia: 5-year follow-up of a randomised trial. Lancet Neurol 2012; 11:1029–1038. S1474-4422(12)70257-0 [pii];10.1016/S1474-4422(12)70257-0 [doi].

[19] Udupa K, Chen R. The mechanisms of action of deep brain stimulation and ideas for the future development. Prog Neurobiol 2015; 133:27–49. S0301-0082(15)00088-X [pii];10.1016/j.pneurobio.2015.08.001 [doi].

[20] Herrington TM, Cheng JJ, Eskandar EN. Mechanisms of deep brain stimulation. J Neurophysiol 2016; 115:19–38. jn.00281.2015 [pii];10.1152/jn.00281.2015 [doi].

[21] Kuhn AA, Kupsch A, Schneider GH, Brown P. Reduction in subthalamic 8-35 Hz oscillatory activity correlates with clinical improvement in Parkinson’s disease. Eur J Neurosci 2006; 23:1956–1960. EJN4717 [pii];10.1111/j.1460-9568.2006.04717.x [doi].

[22] Quinn EJ, Blumenfeld Z, Velisar A, Koop MM, Shreve LA et al. Beta oscillations in freely moving Parkinson’s subjects are attenuated during deep brain stimulation. Mov Disord 2015; 30:1750–1758. 10.1002/mds.26376 [doi].

[23] Ruge D, Tisch S, Hariz MI, Zrinzo L, Bhatia KP et al. Deep brain stimulation effects in dystonia: time course of electrophysiological changes in early treatment. Mov Disord 2011; 26:1913–1921. 10.1002/mds.23731 [doi].

[24] Tisch S, Rothwell JC, Limousin P, Hariz MI, Corcos DM. The physiological effects of pallidal deep brain stimulation in dystonia. IEEE Trans Neural Syst Rehabil Eng 2007; 15:166–172. 10.1109/TNSRE.2007.896994 [doi].

[25] Bocek V, Stetkarova I, Fecikova A, Cejka V, Urgosik D et al. Pallidal stimulation in dystonia affects cortical but not spinal inhibitory mechanisms. J Neurol Sci 2016; 369:19–26. S0022-510X(16)30466-X [pii];10.1016/j.jns.2016.07.053 [doi].

[26] Montgomery EB, Jr. Effects of GPi stimulation on human thalamic neuronal activity. Clin Neurophysiol 2006; 117:2691–2702. S1388-2457(06)01426-X [pii];10.1016/j.clinph.2006.08.011 [doi].

[27] Chiken S, Nambu A. High-frequency pallidal stimulation disrupts information flow through the pallidum by GABAergic inhibition. J Neurosci 2013; 33:2268–2280. 33/6/2268 [pii];10.1523/JNEUROSCI.4144-11.2013 [doi].

[28] Luo F, Kiss ZH. Cholinergic mechanisms of high-frequency stimulation in entopeduncular nucleus. J Neurophysiol 2016; 115:60–67. jn.00269.2015 [pii];10.1152/jn.00269.2015 [doi].

[29] Leblois A, Reese R, Labarre D, Hamann M, Richter A et al. Deep brain stimulation changes basal ganglia output nuclei firing pattern in the dystonic hamster. Neurobiol Dis 2010; 38:288–298. S0969-9961(10)00036-7 [pii];10.1016/j.nbd.2010.01.020 [doi].

[30] Reese R, Charron G, Nadjar A, Aubert I, Thiolat ML et al. High frequency stimulation of the entopeduncular nucleus sets the cortico-basal ganglia network to a new functional state in the dystonic hamster. Neurobiol Dis 2009; 35:399–405. S0969-9961(09)00136-3 [pii];10.1016/j.nbd.2009.05.022 [doi].

[31] Harnack D, Hamann M, Meissner W, Morgenstern R, Kupsch A et al. High-frequency stimulation of the entopeduncular nucleus improves dystonia in dtsz hamsters. Neuroreport 2004; 15:1391–1393. 00001756-200406280-00004 [pii].

[32] Paasonen J, Stenroos P, Salo RA, Kiviniemi V, Grohn O. Functional connectivity under six anesthesia protocols and the awake condition in rat brain. Neuroimage 2018; 172:9–20. S1053-8119(18)30016-8 [pii];10.1016/j.neuroimage.2018.01.014 [doi].

[33] Bode C, Richter F, Sprote C, Brigadski T, Bauer A et al. Altered postnatal maturation of striatal GABAergic interneurons in a phenotypic animal model of dystonia. Exp Neurol 2017; 287:44–53. S0014-4886(16)30344-2 [pii];10.1016/j.expneurol.2016.10.013 [doi].

[34] Gernert M, Richter A, Löscher W. Alterations in spontaneous single unit activity of striatal subdivisions during ontogenesis in mutant dystonic hamsters. Brain Res 1999; 821:277–285.

[35] Gernert M, Hamann M, Bennay M, Löscher W, Richter A. Deficit of Striatal Parvalbumin-Reactive GABergic Interneurons and Decreased Basal Ganglia Output in a Genetic Rodent Model of Idiopathic Paroxysmal Dystonia. The Journal of Neuroscience 2000; 20:7052–7058.

[36] Hamann M, Richter A, Meillasson FV, Nitsch C, Ebert U. Age-related changes in parvalbumin-positive interneurons in the striatum, but not in the sensorimotor cortex in dystonic brains of the dt(sz) mutant hamster. Brain Res 2007; 1150:190–199.

[37] Richter A, Löscher W. Pathophysiology of Idiopathic Dystonia: Findings from Genetic Animal Models. Progress in Neurobiology 1998; 54:633–677.

[38] Richter F, Richter A. Genetic animal models of dystonia: Common features and diversities. Prog Neurobiol 2014. S0301-0082(14)00074-4 [pii];10.1016/j.pneurobio.2014.07.002 [doi].

[39] Avchalumov Y, Volkmann CE, Ruckborn K, Hamann M, Kirschstein T et al. Persistent changes of corticostriatal plasticity in dt(sz) mutant hamsters after age-dependent remission of dystonia. Neuroscience 2013; 250:60–69. S0306-4522(13)00554-X [pii];10.1016/j.neuroscience.2013.06.048 [doi].

[40] Gernert M, Thompson KW, Löscher W, Tobin AJ (2002) Genetically Engineered GABA-Producing Cells Demonstrate Anticonvulsant Effects and Long-Term Transgane Expression when Transplated into the Central Piriform Cortex of Rats. Exp Neurol 176: 183–192.

[41] DeSimone JC, Febo M, Shukla P, Ofori E, Colon-Perez LM et al. In vivo imaging reveals impaired connectivity across cortical and subcortical networks in a mouse model of DYT1 dystonia. Neurobiol Dis 2016; 95:35–45. S0969-9961(16)30163-2 [pii];10.1016/j.nbd.2016.07.005 [doi].

[42] Chiken S, Shashidharan P, Nambu A. Cortically evoked long-lasting inhibition of pallidal neurons in a transgenic mouse model of dystonia. J Neurosci 2008; 28:13967–13977. 28/51/13967 [pii];10.1523/JNEUROSCI.3834-08.2008 [doi].

[43] Paap M, Perl S, Lüttig A, Plocksties F, Niemann C, Timmermann D, Bahls C, van Rienen U, Franz D, Zwar M, Rohde M, Köhling R, Richter A (2020) Deep brain stimulation as a therapeutic option in dystonia: effects of different frequencies in a phenotypic animal model by using an optimized stimulator. Naunyn-Schmiedeberg’s Arch Pharmacol 393: S72.

[44] Morin LPWood RI A stereotaxic atlas of the golden hamster brain. San Diego, London: Academic Press; 2000.

[45] Avchalumov Y, Sander SE, Richter F, Porath K, Hamann M et al. Role of striatal NMDA receptor subunits in a model of paroxysmal dystonia. Exp Neurol 2014. S0014-4886(14)00264-7 [pii];10.1016/j.expneurol.2014.08.012 [doi].

[46] Köhling R, Koch UR, Hamann M, Richter A. Increased excitability in cortico-striatal synaptic pathway in a model of paroxysmal dystonia. Neurobiology of Disease 2004; 16:236–245.

[47] Ashkan K, Rogers P, Bergman H, Ughratdar I. Insights into the mechanisms of deep brain stimulation. Nat Rev Neurol 2017; 13:548–554. nrneurol.2017.105 [pii];10.1038/nrneurol.2017.105 [doi].

[48] Chiken S, Nambu A. Mechanism of Deep Brain Stimulation: Inhibition, Excitation, or Disruption? Neuroscientist 2016; 22:313–322. 1073858415581986 [pii];10.1177/1073858415581986 [doi].

[49] Kirischuk S, Veselovsky N, Grantyn R. Relationship between presynaptic calcium transients and postsynaptic currents at single *gamma*-aminobutyric acid (GABA)ergic boutons. Proc Natl Acad Sci USA 1999; 96:7520–7525.

[50] Dittman JS, Regehr WG. Contributions of calcium-dependent and calcium-independent mechanisms to presynaptic inhibition at a cerebellar synapse. J Neurosci 1996; 16:1623–1633.

[51] Dittman JS, Kreitzer AC, Regehr WG. Interplay between facilitation, depression, and residual calcium at three presynaptic terminals. J Neurosci 2000; 20:1374–1385.

[52] Sciamanna G, Bonsi P, Tassone A, Cuomo D, Tscherter A et al. Impaired striatal D2 receptor function leads to enhanced GABA transmission in a mouse model of DYT1 dystonia. Neurobiol Dis 2009.

[53] Levy LM, Hallett M. Impaired brain GABA in focal dystonia. Ann Neurol 2002; 51:93–101. 10.1002/ana.10073 [pii].

[54] Martella G, Maltese M, Nistico R, Schirinzi T, Madeo G et al. Regional specificity of synaptic plasticity deficits in a knock-in mouse model of DYT1 dystonia. Neurobiol Dis 2014; 65:124–132. S0969-9961(14)00030-8 [pii];10.1016/j.nbd.2014.01.016 [doi].

[55] Scheller U, Lofredi R, van Wijk BCM, Saryyeva A, Krauss JK et al. Pallidal low-frequency activity in dystonia after cessation of long-term deep brain stimulation. Mov Disord 2019; 34:1734–1739. 10.1002/mds.27838 [doi].

[56] Luo F, Kim LH, Magown P, Noor MS, Kiss ZHT. Long-Lasting Electrophysiological After-Effects of High-Frequency Stimulation in the Globus Pallidus: Human and Rodent Slice Studies. J Neurosci 2018; 38:10734–10746. JNEUROSCI.0785-18.2018 [pii];10.1523/JNEUROSCI.0785-18.2018 [doi].

[57] Dostrovsky JO, Levy R, Wu JP, Hutchison WD, Tasker RR et al. Microstimulation-induced inhibition of neuronal firing in human globus pallidus. J Neurophysiol 2000; 84:570–574. 10.1152/jn.2000.84.1.570 [doi].

[58] Hamann M, Richter A. Effects of striatal injections of GABA(A) receptor agonists and antagonists in a genetic animal model of paroxysmal dystonia. Eur J Pharmacol 2002; 443:59–70. S0014299902015467 [pii];10.1016/s0014-2999(02)01546-7 [doi].

[59] Eskow Jaunarajs KL, Bonsi P, Chesselet MF, Standaert DG, Pisani A. Striatal cholinergic dysfunction as a unifying theme in the pathophysiology of dystonia. Prog Neurobiol 2015; 127–128:91–107. S0301-0082(15)00011-8 [pii];10.1016/j.pneurobio.2015.02.002 [doi].

[60] Lavoute C, Weiss M, Rostain JC. The role of NMDA and GABAA receptors in the inhibiting effect of 3 MPa nitrogen on striatal dopamine level. Brain Res 2007; 1176:37–44. S0006-8993(07)01663-0 [pii];10.1016/j.brainres.2007.07.085 [doi].

[61] Liao WT, Tseng CC, Chia WT, Lin CR. High-frequency spinal cord stimulation treatment attenuates the increase in spinal glutamate release and spinal miniature excitatory postsynaptic currents in rats with spared nerve injury-induced neuropathic pain. Brain Res Bull 2020; 164:307–313. S0361-9230(20)30627-4 [pii];10.1016/j.brainresbull.2020.09.005 [doi].

